# Characterising ongoing brain aging and baseline effects from cross-sectional data

**DOI:** 10.1101/2025.01.21.634044

**Authors:** Stephen M. Smith, Karla L. Miller, Thomas E. Nichols

## Abstract

“Brain age delta” is the difference between age estimated from brain imaging data and actual age. Positive delta in adults is normally interpreted as implying that an individual is aging (or has aged) faster than the population norm, an indicator of unhealthy aging. Unfortunately, from cross-sectional (single timepoint) imaging data, it is impossible to know whether a single individual’s positive delta reflects a state of faster ongoing aging, or an unvarying trait (in other words, a “historical baseline effect” in the context of the population being studied). However, for a cross-sectional dataset comprising many individuals, one could attempt to disambiguate varying aging rates from fixed baseline effects. We present a method for doing this, and show that for the common approach of estimating a single delta per subject, baseline effects are likely to dominate. If instead one estimates multiple biologically distinct modes of brain aging, we find that some modes do reflect aging rates varying strongly across subjects. We demonstrate this, and verify our modelling, using longitudinal (two timepoint) data from 4,400 participants in UK Biobank. In addition, whereas previous work found incompatibility between cross-sectional and longitudinal brain aging, we show that careful data processing does show consistency between cross-sectional and longitudinal results.

## 1 Introduction

Researchers have proposed the concept of “brain age” as a way of deriving markers of neurobiological aging, based, for example, on imaging data (Franke et al., 2010; Cole and Franke, 2017; Cole et al., 2017). This typically proceeds by using a dataset with many subjects’ imaging data, modelling how the average brain changes with aging, and then using this to estimate each individual’s apparent brain age. The difference between brain age and actual age is used to infer whether the subject’s brain appears to have aged more or less than the population norm for that true age (Figure 1A). This difference is often referred to as “brain age delta” or “brain age gap estimate” (BrainAGE, Franke et al., 2010). Individual subjects will of course follow their own time-varying aging trajectory, with external influences on aging changing over time; however, here, we are focussed on what is estimable in terms of the overall (average) population patterns of aging. Biological age modelling is not of course limited to the brain; see Moqri et al., 2023 for a thorough review of more general considerations for various organs and modelling approaches.

**Figure 1:**
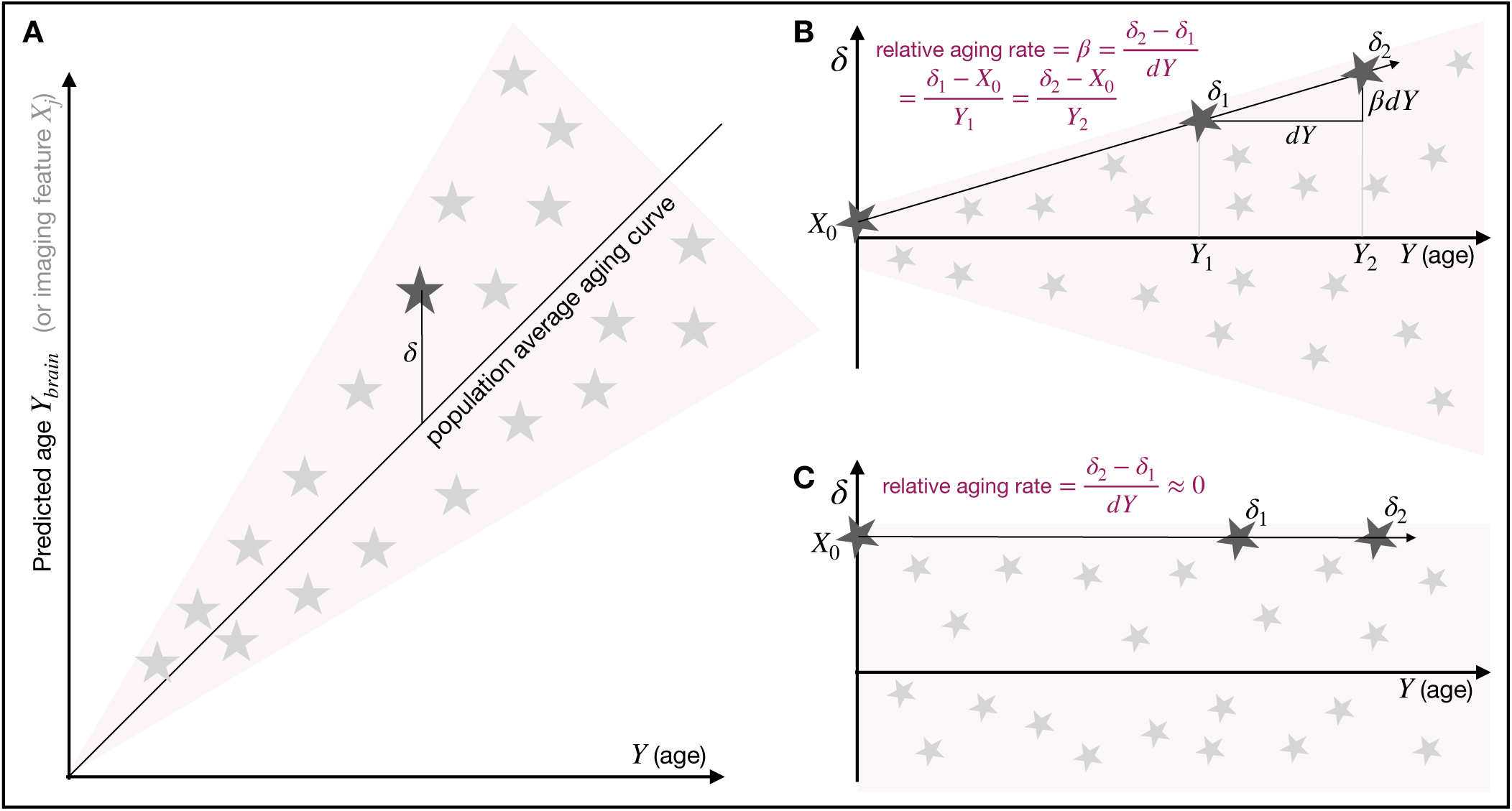
Illustrations of the relationships between age, predicted age, and brain age delta. A. Relationships before subtracting the population average aging curve. B. Delta (after subtracting the average curve) in the scenario where each subject has a different aging rate. A subject’s aging slope consistently describes both the rise in delta from its baseline value *X*_0_, and the longitudinal change in delta between two timepoints. C. Delta in the case where there are no subject variations in ongoing aging rate, and therefore delta is not changing over time; hence there cannot be any relationship between longitudinal and cross-sectional delta.

A common interpretation of brain age delta is that positive brain age delta means accelerated brain aging. If different subjects are truly aging at different rates, we should see that the average magnitude of delta (the between-subject spread of delta values) is changing with increasing age (Figure 1B). We refer to this as “delta-expansion”. If we do not observe expansion, the implication is that delta is not systematically changing with age, and that delta cannot be interpreted as reflecting an ongoing aging process. In this case, it must instead reflect intrinsic state effects present from a younger age (i.e., “baseline” differences, Figure 1C). Put more simply, learning a spatial pattern in the brain which reflects aging on average in the population does not guarantee that the pattern seen in any individual can safely be interpreted as purely reflecting aging; it is also possible that the pattern reflects other sources of between-subject variation. For example, total grey matter volume is a good marker of aging, but is also highly variable between subjects of different overall body size.

In S. Smith et al., 2019 we had proposed a simple measure reflecting the amount of delta-expansion, finding a very small amount of expansion when using 2,641 cross-modal IDPs (imaging-derived phenotypes, Miller et al., 2016) from 19,000 subjects in UK Biobank; the expansion only amounted to 0.4y over the 45-80y age range. Here, we propose a more precise measure of delta-expansion, which splits the total population delta variation into the variance explained by baseline differences vs. the variance explained by delta-expansion. The latter indicates how confidently we can interpret brain-age-delta values as reflecting an ongoing brain aging process.

Vidal-Pineiro et al., 2021 investigated empirically whether brain age delta reflects between-subject variability in ongoing aging vs. baseline effects, noting that the “assumption that brain age delta reflects an ongoing process of faster or slower neurobiological aging implies that there should be a relationship between cross-sectional and longitudinal estimates of brain age” (a point illustrated in Figure 1B). The study found “no association between cross-sectional brain age and the rate of brain change measured longitudinally”, and that “Birth weight was significantly related to brain age delta”. The study concludes that “cross-sectional brain age has low validity as an index of [between-subject variations in the rate of] brain aging”.

Traditional methods estimate a single (“all-in-one”) brain-age-delta per subject, using all imaging data available. This approach is likely to pool across multiple biologically-distinct modes of aging, which would make meaningful interpretation harder. We previously showed that different modes of brain aging have different average aging curves, distinct biological characteristics (distinct sets of genetic and phenotypic associations), and different amounts of delta-expansion (S. M. Smith et al., 2020). Different modes of aging were found to have rich and distinct genetic associations, whereas, by implicitly combining these modes together, the all-in-one brain age model diluted their unique neurobiological associations, giving a much weaker and less rich set of genetic associations. This previous work combined across structural, functional and diffusion imaging modalities, but this is not essential for deriving multiple modes of aging. Even if only one modality is available, (e.g., just T1-weighted structural imaging or functional connectivity matrices), there may be several separate modes of aging, reflecting distinct changes in tissue volumes, lesions, or tissue intensities, and engaging different parts of the brain (Kessler et al., 2016; Vinke et al., 2018).

The above considerations (and modelling approaches) are relevant for cross-sectional data (single timepoint per subject), which is important because most imaging studies are cross-sectional. Here, we apply our modelling cross-sectionally to 4,400 UK Biobank subjects that also have a second imaging timepoint (time between scans 2.6 ± 1.0 years). This enables us to verify that our modelling results are consistent with the measured longitudinal changes; crucially, this consistency is only seen when applying careful data processing to avoid specific modelling pitfalls. We verify that some modes of brain aging reflecting strong delta-expansion are correctly interpreted, by showing congruence with longitudinal results. We find a strong correlation between delta-expansion and the consistency between *δ*_cross_ (the average of the two timepoints’ separate estimates of delta) and *δ*_long_ (the difference between the two estimates).

We also revisit the distinction between all-in-one models vs. multiple modes; contrasting examples are shown in Figure 2, using data from 4,400 subjects in UK Biobank. In (A), a standard all-in-one model of brain aging shows good age prediction, but little delta-expansion, and hence should not be interpreted as predominantly representing ongoing aging. In (B), the mode of brain aging that has the strongest age correlation shows no delta-expansion. In (C), another mode shows clear delta-expansion, hence likely reflecting ongoing aging. More quantitatively, the all-in-one delta reflects only small amounts of delta-expansion, with the spreading accounting for just 23% of the population variability in delta, compared with the individual modes’ spreading accounting for up to 68%.

**Figure 2:**
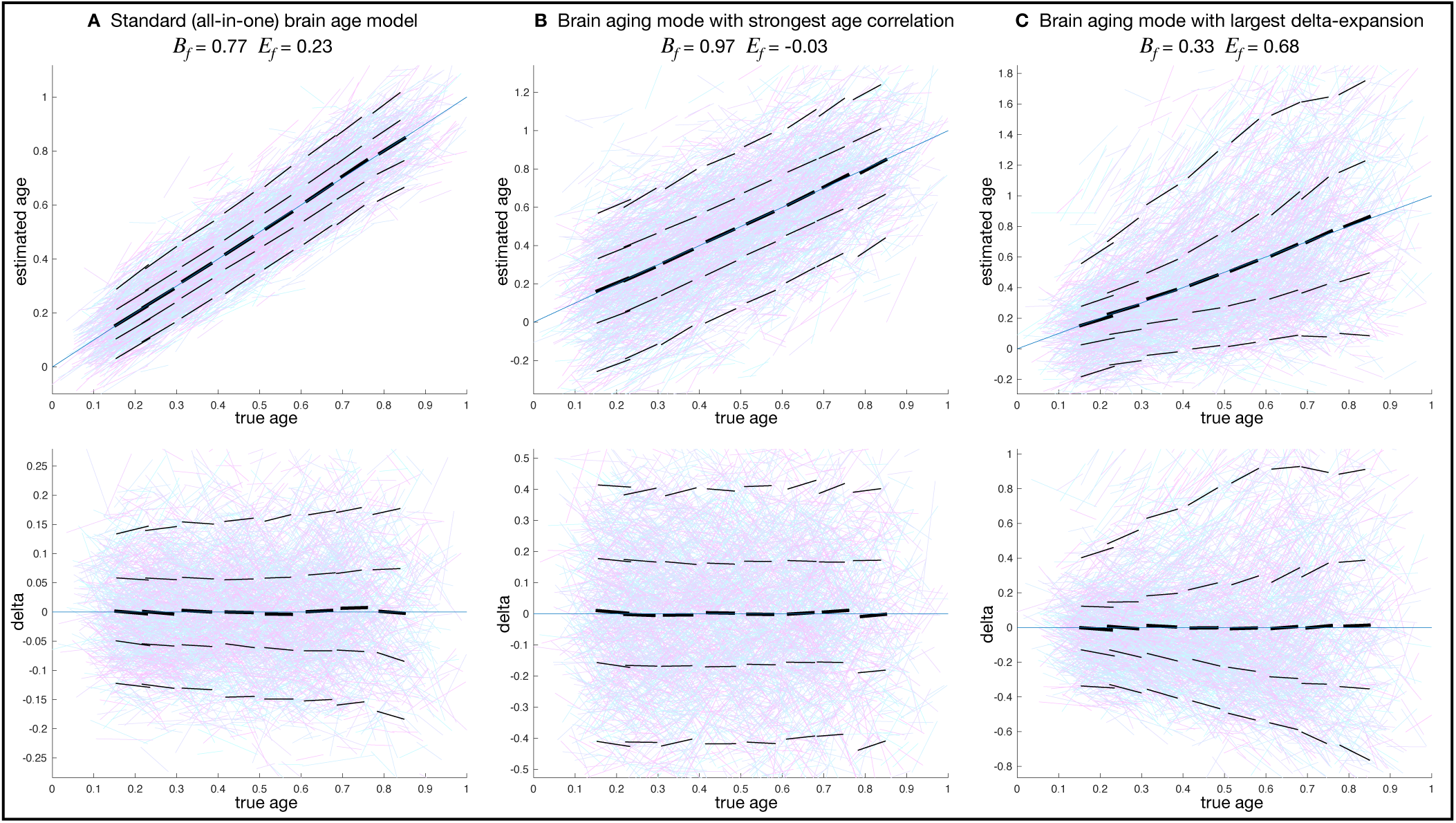
Predicted age (above) and brain age delta (below) vs. true age. Age is normalised into the range 0–1. Data are visualised as lines between two imaging timepoints, either representing local population averages (black) or individual subjects (coloured). The two timepoints from each subject are modelled separately, meaning that the age modelling here is purely cross-sectional. However, for visualisation purposes, the coloured line segments show the age estimation for the two timepoints of each subject. The thick black lines are similar, but averaged over all subjects within a nearby time-window (having a mean age within 0.1 of the window centre). The thinner black lines show similar windowed averages, but for the median (and 90th percentile) subject with positive delta, and median (and 10th percentile) subject with negative delta. *E_f_* is the fractional population variance explained by delta-expansion, and *B_f_* is the fractional variance explained by baseline effects. **A.** The all-in-one age prediction (using all 3,939 imaging-derived phenotypes from 6 structural and functional imaging modalities). **B.** The mode of brain aging showing strongest correlation with age. **C.** The mode of brain aging showing maximal delta-expansion with increasing age. The all-in-one brain aging model and aging mode 1 show almost no visually-apparent delta expansion; indeed, in (B), *E_f_* is reported as negative, because the best-fitting model was delta-contraction (not expansion).

Brain age deltas are estimated at the subject level for all methods discussed here, and any downstream analyses using the deltas (including group differences or correlations with non-imaging variables) would proceed as normal (albeit with one set of deltas per mode in the case of multiple modes of aging). The specific focus here is on the interpretation of deltas, and on evaluating whether delta expansion is more likely to be found when estimating multiple modes.

## 2 Methods

### 2.1 Brain age models

We first describe a typical framework for brain age and brain age delta estimation, and then describe how delta-expansion (with increasing age) can be modelled. Although some approaches to brain age estimation use nonlinear methods, the following linear formulation directly translates to nonlinear prediction, because it does not matter how the brain age (and the delta) is estimated, in order for delta-expansion estimation to be applied.

In Figure 1, we visualise the general properties of age-dependent imaging features in the context of attempting to distinguish between baseline effects and delta-expansion. The vertical axis in Figure 1A can represent either predicted age (in the context of all-in-one modelling) or an age-dependent image-based feature. There are many ways to derive or identify such image features. The traditional brain age approach explicitly fits the imaging data, in the form of voxels or IDPs, to age. This fit produces a single (all-in-one) imaging feature that is typically interpreted at the subject level as brain age. However, one does not need to explicitly fit to age to identify features that can be meaningfully analysed this way. In (S. M. Smith et al., 2020) we used a different approach to derive multiple independent modes of brain aging. We used a data-driven analysis (independent component analysis) to cluster imaging data that has been pre-processed to enhance aging effects. Some of these modes (ICA components) exhibit strong age dependence and can thus be considered as age-relevant imaging features. In the simplest case, one could consider age-dependent IDPs or even individual voxels in this context, although, as discussed at the end of this section, this is unlikely to be useful.

Prediction of age (*Y*) is, in the case of linear methods, carried out using a matrix of imaging features *X* as predictors. The columns of *X* could be imaging-derived phenotypes (IDPs), or voxels, or “meta-IDPs”, for example, by dimensionality reduction or feature selection from the original matrix of IDPs or voxels. *Y* and *X* contain one entry (row) per subject, and for convenience we rescale *Y* to lie in the range 0–1 (with *Y* = 0 corresponding to the youngest subject in the dataset).

The simple linear all-in-one model (Figure 1A) for brain age prediction is typically:

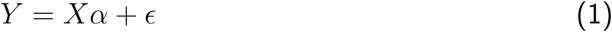

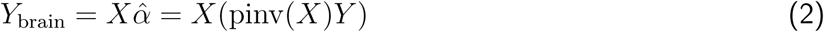

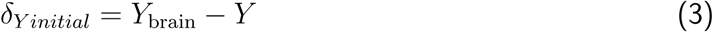

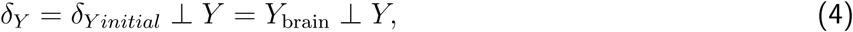

where *α* is the vector of age-prediction regression coefficients, *Y*_brain_ is predicted age (brain age) and ⊥ *Y* means orthogonalise with respect to *Y*, i.e., we regress *Y* out of the initial delta (*δ_Y initial_*) to avoid underfitting bias (S. Smith et al., 2019).

As an alternative formulation, we can switch from viewing age as the dependent variable to having aging as the causal factor affecting imaging features (which are therefore modelled as a function of age). Instead of the standard all-in-one formulation above, we consider each imaging feature to follow a simple generative model that begins with the population average age dependence, and then adds each individual’s baseline and age-dependent deviation from that population average (Figure 1B). We now describe the modelling for a single imaging feature *x*, which for feature *j* is column *j* in *X*. For subject *i* (the *i*th value in *x*) we have:

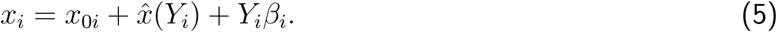

The population average aging curve is captured by *x*^*_i_*, here a quadratic function of age *Y_i_* (a more complicated form can be used where the data supports and requires this). Two subject-specific terms are added on top of the population average. First, the subject’s age-independent baseline (the value it would have at age *Y* = 0) is given by *x*_0_*_i_*. Second, the subject’s aging rate *β_i_* is multiplied by their age to capture faster/slower aging relative to the population average; this model assumes that this rate is not changing over time. By definition both *x*_0_ and *β* are zero mean (i.e., when averaged across all subjects).

Importantly, because we have cross-sectional data (a single datapoint per subject), we cannot estimate the baseline and aging rates at a single subject level (i.e., we cannot estimate *x*_0_*_i_* and *β_i_*). Nevertheless, this generative model includes both effects, which we need to separate at the population level if we wish to distinguish true aging effects from baseline.

Subtracting the population mean aging curve *x*^*_i_* from *x_i_* gives the delta,

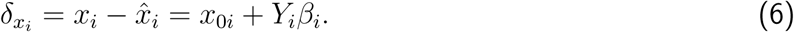

Although we cannot estimate *x*_0_*_i_* and *β_i_* for any given subject (we cannot estimate their distinct effects on a single datapoint), from the age-varying changes seen across the population, we can estimate their variances:

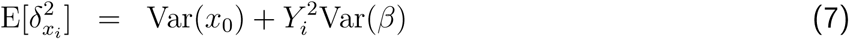

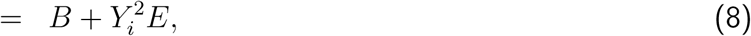

where *B* (“baseline” subject variability Var(*x*_0_)) and *E* (“expansion” subject variability Var(*β*)) are the unknown scalar quantities to be estimated, and we have assumed that *x*_0_ and *β* are uncorrelated, an issue we revisit below (Section 3.3). Note that we are not reducing *δ_xi_* to a single variance, but have an expected value for *δ*^2^*_xi_* that is age-dependent; we then fit this model using a maximum-likelihood iterative optimiser, initialised by a methods-of-moments fit (see Appendix and code linked below). This is one particular type of heteroscedastic model induced by our particular modelling assumptions; alternatively, a Generalised Additive Model (Hastie & Tibshirani, 2017) could be used, but would lack the same interpretability. Considering the total average variability

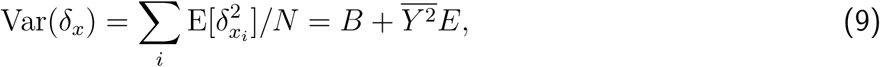

where *N* is the sample size and *Y* ^2^ = ∑*Y* ^2^*/N*, we can write the fractional baseline variance *B_f_* = *B/*Var(*δ_x_*) and fractional expansion variance *E_f_* = *Y* ^2^*E/*Var(*δ_x_*). Note that *B_f_* + *E_f_* = 1. We estimate the mean age curve and *B*, *E*, *B_f_* and *E_f_* for each imaging feature (and for the all-in-one model).

The above assumes that if delta-expansion is apparent in the data, it is more likely to be positive (larger delta magnitudes with increasing age) than negative (reducing delta). Empirically we do indeed find this to be the case. However, in order to allow for the possibility of contraction (negative expansion), we fit the expansion model twice, once as written above, and once after replacing *Y* with 1 − *Y* in equation 8. The likelihoods of the two model fits are then compared, and the best-fitting model used. In the case of contraction, the “baseline” variability relates to the cross-subject spread at the oldest age of the sample, not the youngest. The model also makes various other simplifying assumptions. For example, subject-specific aging rate may not be constant over time; the expansion could be quadratic, or varying across subjects in an even more complex way, including different subjects having different ages at which the aging rate changes. However, for many cross-sectional datasets, more complex modelling is unlikely to be identifiable (supported by the data), and the estimation of *E_f_* as a first-order indicator of expansion is hopefully useful.

The generative model is fit separately for each imaging feature *j* in *X*. We believe this is a good approach when different imaging features reflect distinct aging processes (and hence distinct dependence on age). On the other hand, if the imaging features are voxels or IDPs, it is likely to be overwhelming to interpret, and noisy, to model each feature in isolation. We therefore recommend pre-clustering voxels or IDPs into a smaller number of modes of brain aging (S. M. Smith et al., 2020), for example, using independent component analysis. This results in a more compact, higher signal-to-noise-ratio, version of *X*. For any new imaging feature (final column in *X*), because the original features (IDPs or voxels) have only been clustered together on the basis of having a similar aging curve to each other, the intepretability of the individual modes does not suffer.

### 2.2 Real data

Our real data evaluations used IDPs from 4,400 subjects (54% female) in UK Biobank, using 6 imaging modalities (T1, T2-FLAIR, swMRI, dMRI, rfMRI, task-fMRI, as described in Miller et al., 2016). The mean age at first imaging scan was 62.1y (standard deviation 7.4y, range 46.1-80.8y). The mean interscan interval was 2.6y (standard deviation 1.0y, range 1.0-7.0y). UK Biobank has approval from the North West Multi-centre Research Ethics Committee (MREC) to obtain and disseminate data and samples from the participants (http://www.ukbiobank.ac.uk/ethics/), and these ethical regulations cover the work in this study. Written informed consent was obtained from all participants.

We used 3,939 IDPs spanning a range of structural, diffusion and fMRI phenotypes. These are estimated by our analysis pipeline developed and run on behalf of UKB (Alfaro-Almagro et al., 2018). For each IDP, outlier values were removed, where absolute deviation from the median was greater than 5 times the median absolute deviation from the median. The following confounds were regressed out of the data: sex, head size, two measures of scanner table position, three measures of head motion estimated from rfMRI, scanning site, eddy-correction method, and an eddy current QC metric; for full details of confounds used, see the analysis code linked below. We used data from all subjects currently available (as of summer 2024) with two timepoints, and where less than 2% of IDPs were missing in both timepoints. The two timepoints for each subject were included as separate samples, meaning that the IDP matrix had 8,800 rows. This is important (see below), to ensure that the cross-sectional analyses are not biased towards either timepoint.

Following our previous approach for estimating multiple modes of brain aging (S. M. Smith et al., 2020, where the choice of the number of modes was discussed and investigated in depth), we first reduced the IDP matrix from 3,939 IDPs to 16 components using singular value decomposition (SVD) while simultaneously imputing missing data using SVD-based imputation with soft eigenvalue shrinkage (https://fsl.fmrib.ox.ac.uk/fsl/docs/#/resting_state/fslnets). Similar to that previous work, in order to help focus this data-driven decomposition on age-related population modes, each IDP was rescaled by the square of the correlation between the IDP and age, before applying the SVD (noting that we previously demonstrated that this step does not affect the validity of the age associations of the final modes). The 16 horizontal (IDP weights) eigenvectors from the SVD were then fed into the FastICA (Hyvärinen, 1999) algorithm for independent component analysis, to find independent sources in the IDP (not subject) direction.

This estimation of multiple modes (dimensionality reduction followed by ICA unmixing) can simply be thought of as a soft clustering, grouping multiple IDPs together where they have similar variation across subjects. This reduction (from voxels or IDPs to modes) can easily be translated from one study to another if they have a compatible set of imaging modalities; just as a trained all-in-one model might hope to translate from one study to another, it would also be the case that a multiple-modes decomposition would translate in the same way (by carrying across the spatial basis sets from the SVD/ICA).

ICA produced 16 components, each described by a vector of 3,939 IDP weights reflecting the strength of association of each IDP with that component. After projecting the ICA mixing matrix back to the original data dimensions, each component has an accompanying vector of 2 × *N_subjects_* weights, describing the strength of each component for a given subject-timepoint (which could be interpreted as their “brain age” in that component). Enforcing independence (via ICA) in the IDP direction does not place any restriction on the correlation between the subject-weight vectors, apart from requiring the matrix of these to be full rank. For convenience of reporting (and without quantitatively altering the modelling) the subject-weight vectors were sign-flipped so as to positively correlate with age, and sorted into decreasing strength of correlation with age; matching changes were made to the IDP weights vectors. At this point the 16 subject-weight vectors comprise our image feature matrix *X*, i.e., are the distinct modes of brain aging.

For the all-in-one model of age prediction, we used all of *X* (i.e., all imaging features after the ICA reduction) in a simple multiple regression, as described above. As found previously (S. Smith et al., 2019), this gives better age prediction than using the original full set of thousands of raw IDPs as predictors. Separately, for investigating multiple modes of brain aging, each of the 16 columns in *X* was modelled as above to give a separate set of deltas for each mode. For each set of delta values, we applied the delta-expansion modelling, resulting in estimation of *B* and *E* (and *B_f_* and *E_f_*) for each set. The choice of SVD/ICA dimensionality did not have much effect on the all-in-one modelling; for example, *E_f_* only had a 2.5% coefficient-of-variation from 14 dimensions up to 30. Similarly, for multiple modes of aging also, the range of delta-expansion values did not vary greatly with dimensionality.

For the investigations here, where we have two timepoints for each subject, we pooled all data into a single “cross-sectional” analysis, resulting in a single set of 2*N_subjects_* all-in-one delta values, and 16 individual modes’ 2*N_subjects_* delta values. This allows us to then investigate relationships between cross-sectional and longitudinal deltas in an unbiased way (see the following paragraph). Pooling the data in this way does not significantly change the cross-sectional modelling; for example, the regression coefficients from the all-in-one brain age prediction correlate almost perfectly (*r* = 0.98) between using only time1 vs. time2 data, and those two sets of coefficients both correlate with the 2*N_subjects_* all-data coefficients with *r* = 0.99. Similarly, the estimated baseline and delta-expansion factors were almost identical (with vs. without pooling); fitting paired data with the cross-sectional model makes the standard errors overly optimistic, so we inflate the reported real-data SEs by a factor of √2 as a conservative adjustment. Importantly, we only pool the two timepoints to investigate the validity of the delta-expansion estimates; in a conventional cross-sectional study, one would simply use all of the single-timepoint data available when carrying out the analyses proposed in this work.

Finally, from each set of delta values, we separated the vector into the two timepoints, so that we could then compute the average of the two deltas (forming the cross-sectional mean *δ*_cross_), and also the difference normalised by the between-scan interval (to find the longitudinal change *δ*_long_). In cases of delta-expansion, we would hope to find consistency between these, as evaluated by correlating *δ*_cross_ with *δ*_long_. For such evaluations it is vital to correlate *δ*_long_ with the average of the two timepoints’ deltas, and not just the first (or the second) timepoint, in order to avoid strong noise-driven negative (or positive) bias in the correlations.

### 2.3 Simulations

The primary goal of our simulations was to take advantage of knowing the ground-truth values underlying the simulation to evaluate aspects of the modelling that cannot be directly verified from real data. The most important aspects are the subject-level baseline and age scaling rate. Our modelling cannot estimate these at the subject level, but, because we know their ground truth values, we can evaluate the extent to which estimated subject-level deltas correlate with (the ground truth values of) these two factors.

Secondly, we can evaluate how identifiable the models are; just because a given generative model is used to simulate data, that does not mean that its parameters are necessarily identifiable when fitting the same model to the simulated data. We primarily were concerned with ability to recover *E*, *B*, *E_f_* and *B_f_* .

In order to test the different brain aging models, we generated simulated data following the same generative models as described above; specifically, we use Eq. 5 to generate the data for each brain aging mode, having chosen suitable values for the free parameters. The simulation scripts (including controlling parameters) are linked below. We generated 16 aging modes following Eq. 5, with the addition of separate measurement noise. This noise would be indistinguishable from *x*_0_ for single-timepoint data, but we generate two timepoints for each subject, matching our real data. Hence *x*_0_ is consistent between the two timepoints, whereas the added measurement noise is independent. On average, 10% of modes are set for delta-contraction, with a 5-fold reduction in the strength of the contraction (compared with the delta-expansion for expanding modes). As the modes of aging are not constrained (by the ICA) to be orthogonal, in order to generate realistic levels of between-mode correlations (in both modes and deltas), the subject-specific aging rates and baseline effects were simulated from a multivariate normal with controlled amounts of positive correlation matching that seen in the real data. Second timepoint data was generated with a time interval of 0.05 (compared with the full age range of 0:1).

The simulation parameters (data dimensions, and the distributions of population average aging slopes, baseline delta effects, aging-rate variation, added noise, and correlations between modes) were chosen to result in simulated data having identical characteristics to the real data. The simulation was run 100 times, so that we could report mean and standard deviation accuracies across multiple realisations of the data.

## 3 Results

### 3.1 Real data

The all-in-one modelling gave high age prediction accuracy: correlation between true and predicted age was *r*= 0.91, and mean absolute delta was 2.65y. From the multiple modes modelling, most modes correlated reasonably strongly with age, with the first mode’s correlation being *r* = 0.55, and all except one correlating with age with *r >* 0.3 (Figure 3F, plot 1).

**Figure 3:**
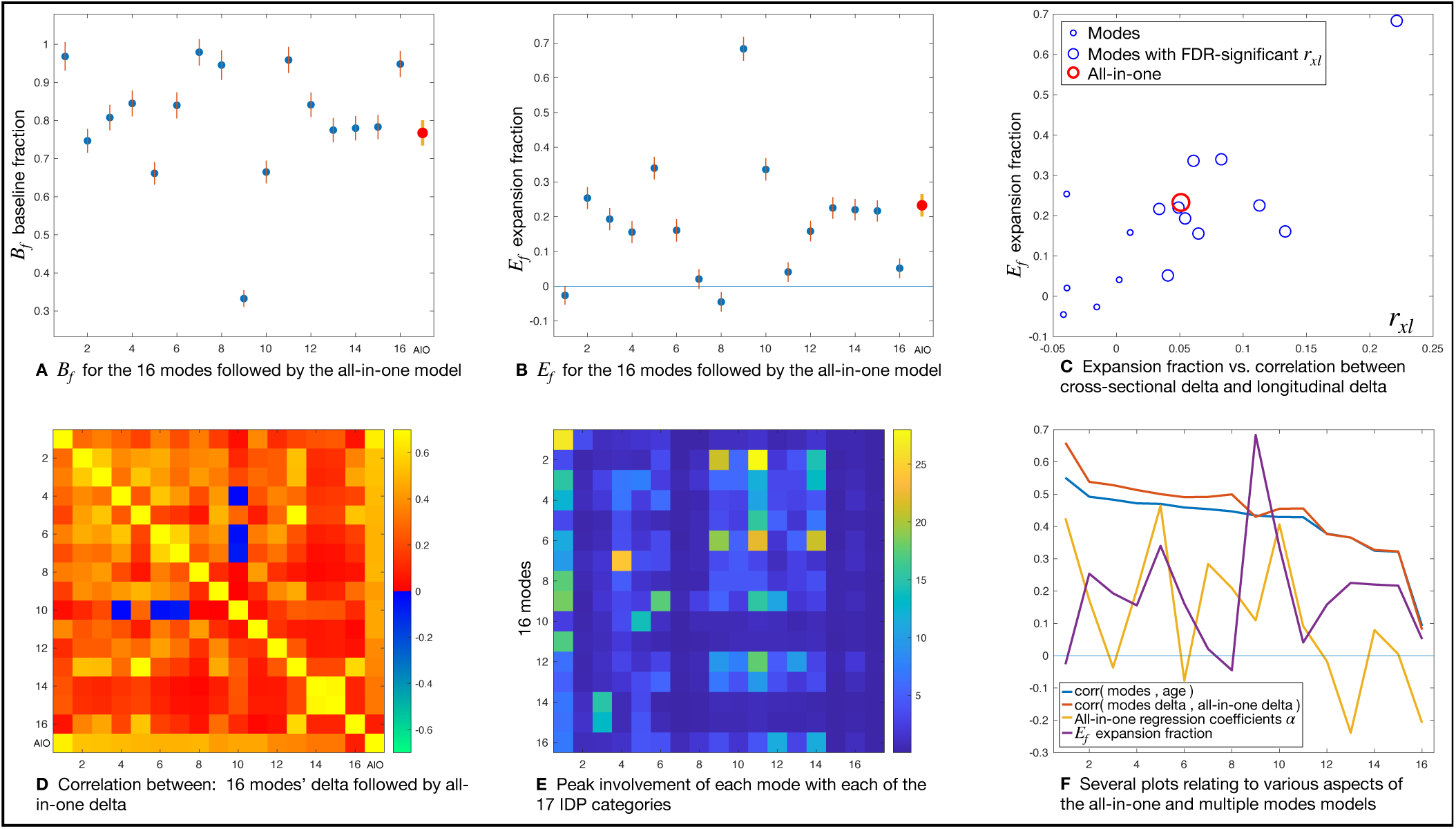
Detailed results for the all-in-one brain aging model and the 16 separate aging modes; see main text for further details. A,B. The baseline and expansion fractions for the 16 modes and the all-in-one model. C. The relationship between delta-expansion and the correlation *r_xl_* between *δ_cross_* and *δ_long_*. D. The correlations between delta of all modes with each other. E. How the 16 brain aging modes map onto different categories of IDPs. The 17 IDP categories on the x-axis are: 1. regional and tissue volume; 2. cortical area; 3. cortical thickness; 4. regional and tissue intensity; 5. cortical grey-white contrast; 6. white matter hyperintensity volumes; 7. regional T2*; 8. regional QSM (quantitative susceptibility modelling); 9. WM (white matter) tract fractional anistropy; 10. WM tract mode; 11. WM tract diffusivity; 12. WM tract intra-cellular volume fraction; 13. WM tract orientation dispersion ; 14. WM tract isotropic or free water volume fraction; 15. tfMRI (task functional MRI) activation; 16. rfMRI (resting fMRI) node amplitude; 17. rfMRI connectivity. F. Plots relating to various aspects (see legend) of the modelling.

However, the all-in-one model performed badly from the point-of-view of displaying delta-expansion, with a modest *E_f_* = 0.23, and hence baseline effects explaining 77% of the between-subject variance (*B_f_* = 0.77). This is visualised in Figure 2A, showing only very weak delta-expansion. Correlation between *δ*_cross_ and *δ*_long_ was low but positive and statistically significant (*r* = 0.05*, p* = 7.5 × 10^−4^). Although in other contexts it might be argued that explaining 23% of the variability in a brain imaging measure is actually rather impressive, here we need to keep in mind that delta estimated for any given subject cannot be unambiguously ascribed to reflecting rate of ongoing aging, and therefore this result is saying that on average the variance in brain age delta is three times more strongly explained by fixed baseline effects than by variations in rate of aging.

The multiple modes of brain aging showed a range of delta-expansion values, the largest being mode 9, with high expansion *E_f_* = 0.68 (Figure 2C) and significant correlation between *δ*_cross_ and *δ*_long_ (*r* = 0.22*, p* = 6.0 × 10^−50^). Notably, the mode with the strongest age correlation (mode 1) had close to zero delta-expansion (Figure 2B).

More details are shown in Figure 3. 3A and B show the baseline and expansion fractions for the 16 modes and the all-in-one model. Where the expansion modelling has estimated negative delta-expansion (i.e., contraction of the average delta magnitude with increasing age), *E_f_* is shown as being negative. The maximum-likelihood model fitting produces estimates of standard errors on *B* and *E* (Appendix A); these are indicated by the vertical lines. A wide range of expansion fractions is found.

Figure 3C shows the relationship between delta-expansion and the correlation *r_xl_* between *δ*_cross_ and *δ*_long_. Recalling Figure 1B and C, *r_xl_* captures whether the cross-sectional delta is actually getting larger over time. If there is truly an aging expansion effect, then *δ*_cross_ should be predictive of the change in delta between timepoints, *δ*_long_, whereas, if there are only baseline effects, then *δ*_cross_ is not age dependent and *δ*_long_ will simply reflect measurement noise. One would therefore hope that *E_f_* and *r_xl_* are directly linked; we only expect a (strong and positive) correlation between cross-sectional and longitudinal delta in the scenario of delta-expansion, and this is indeed what we find here, with the correlation across modes *r*(*E_f_, r_xl_*) = 0.77. The *r_xl_* correlations are limited in strength (maximum of *r_xl_* = 0.22) primarily because *δ*_long_ is intrinsically noisy, being estimated via the subtraction of the two timepoints’ deltas; however, it is notable that the correlations are mostly positive and FDR-significant (10 of 16 modes).

Figure 3D shows the correlations between deltas of all modes with each other. The modes themselves are generally expected to correlate positively with each other, because they are all oriented to correlate positively with age, and for most modes this correlation is reasonably strong. However, there is no equivalent reason why the modes’ deltas should all correlate positively with each other (that is, after regressing out age trends and imaging confounds). The between-mode correlations (correlations between columns of *X*) are all positive (*r* = 0.39 ± 0.16), and the between-mode delta correlations are almost all positive (*r* = 0.24 ± 0.17, with just 3 negative correlations, the most negative being *r* = −0.07). The dominantly positive correlations between modes’ deltas is presumably due to some sharing of (baseline and/or expansion) causal factors. (This result could of course be different if a different method was used for identifying distinct modes; for example, using PCA or subject-direction-ICA would result in orthogonal modes and probably largely-orthogonal deltas.) (D) also shows correlations of all modes’ deltas with the all-in-one delta; as expected (Appendix B), these correlations are all positive, reinforcing the interpretation of all-in-one delta as being a munged combination of the modes’ deltas.

Figure 3E illustrates how the 16 brain aging modes map onto different categories of IDPs (with Table 1 listing the main associations for individual IDPs within these categories). For each mode, the ICA source weights (one value per IDP) are used to display the largest weight (corresponding to a single IDP) within each category. Modes of aging having strong age correlation but very low delta-expansion (1, 7, 8, 11) are dominated by regional and tissue volumes and intensity. Modes showing strong delta-expansion involve a wider range of IDP categories, most notably diffusion MRI metrics, white matter hyperintensities (lesions), grey-white contrast and cortical thickness.

**Table 1:** Lists of IDPs and non-neuroimaging measures most strongly associated with all of the aging models. For non-imaging measures, the strongest classes of associations above |*Z| >* 4 are listed. Abbreviations: AIO all-in-one brain age model; GM grey matter; MD mean diffusivity; T1 T1-weighted structural MRI; FA fractional anisotropy; MO tensor mode; WM white matter; ISOVF isotropic volume fraction; L1 MD along dominant tract direction; CSF cerebrospinal fluid; ICVF intracellular volume fraction; ECG electrocardiogram; BP blood pressure; BMI body mass index; PRS polygenic risk score.

| | $E_f$ | $r_{\Delta}$ | IDPs | non-neuroimaging measures |
| --- | --- | --- | --- | --- |
| AIO | 0.23 | 0.05 | volume of GM, MD in in anterior thalamic radiation and anterior internal capsule, T1 contrast across cortical grey-white boundary, T1 intensity in 3rd ventricle | ECG intervals, heart rate, BP, smoking history, lung capacity, overall health rating |
| 1 | 0.03 | -0.02 | volume of GM | bone mineral density, smoking history, ECG intervals, cognitive trail making speed |
| 2 | 0.25 | -0.04 | FA/MO/MD in fornix | thigh muscle fat, cognitive symbol digit matches, vascular/heart problems |
| 3 | 0.19 | 0.05 | volume of brain (GM+WM), MD/ISOVF in corpus callosum | BP, cognitive reaction time, bone mineral density, BMI, myocardial wall thickness |
| 4 | 0.16 | 0.06 | volume of nucleus accumbens, L1 in anterior thalamic radiation | bone mineral density |
| 5 | 0.34 | 0.08 | MD in in anterior thalamic radiation | BP, heart rate, cholesterol, myocardial wall thickness |
| 6 | 0.16 | 0.13 | volume of CSF, FA/MD in fornix | cognitive symbol digit substitution speed, hand grip strength |
| 7 | 0.02 | -0.04 | T1 intensity in CSF, choroid plexus and thalamus | body size, bone mineral content |
| 8 | 0.05 | -0.04 | volume of thalamus | bone mineral density |
| 9 | 0.68 | 0.22 | volume of WM hyperintensities, MD in superior fronto-occipital fasciculus | diastolic BP, heart rate, BMI, PRS for ischaemic stroke, myocardial wall thickness |
| 10 | 0.34 | 0.06 | T1 contrast across cortical grey-white boundary | bone mineral density, fatigue |
| 11 | 0.04 | 0.00 | volume of hippocampus | bone mineral density |
| 12 | 0.16 | 0.01 | L1 in anterior internal capsule and anterior and superior thalamic radiations | heart rate |
| 13 | 0.23 | 0.11 | MD in posterior corona radiata and posterior thalamic radiation | BP |
| 14 | 0.22 | 0.05 | thickness of cortex, left hemisphere | ECG load during exercise |
| 15 | 0.22 | 0.03 | thickness of cortex, right hemisphere | creatinine (kidney function), ECG load during exercise |
| 16 | 0.05 | 0.04 | ICVF and ISOVF in many WM tracts | aorta area, bone mineral density |

Figure 3F shows several final plots relating to various aspects of the modelling. The first two plots show the correlations between the brain aging modes and age, and the correlation between the modes’ deltas and the all-in-one deltas. The two plots are similar, which is generally to be expected (Appendix B), and with strongly positive correlations (for all except the last mode). The third plot shows the all-in-one model regression coefficients *α*. Despite each *X_j_* being oriented to correlate positively with age, not all the regression coefficients are positive. This could arise for several reasons, for example, different imaging features (or modes, *X_j_*) needing to partially cancel each other out because they contain non-age-related structured effects reflecting other aspects of brain biology (see Discussion); however, the negative values could also simply be caused by having highly correlated regressors. The final plot shows the fractional delta-expansion values *E_f_* ; these were already shown in panel (B), but are included here to illustrate that there is no simple link between delta-expansion and other measures such as strength of age correlation.

Table 1 lists the IDPs most strongly associated with each of the brain aging models/modes (according to the multiple regression coefficients for the all-in-one model, and the ICA coefficients for the multiple modes of aging). It also lists the non-neuroimaging variables most strongly associated with each corresponding set of deltas. The individual modes are generally associated with focussed sets of IDPs, whereas the all-in-one model is associated with a larger, less focussed set of IDPs. The mode of brain aging showing the strongest correlation with age (mode 1, as visualised in Figure 2B) primarily relates to the total volume of grey-matter, and for this mode the most strongly associated non-imaging variables are mainly other biological measures also affected by aging (e.g., bone density, Cordes et al., 2016; Loskutova et al., 2009), with just smoking history relating to likely root causes of advanced or unhealthy aging. Given that there is no delta-expansion associated with this mode, one should conclude that the causes of subject variability in this mode relate more to earlier-life effects rather than differences in ongoing aging rate. The mode of brain aging showing maximal delta-expansion with increasing age (mode 9, as visualised in Figure 2C) primarily relates to the total volume of white-matter hyperintensities (“lesions”) and diffusivity in some major white matter tracts such as the fronto-occipital fasciculus; the non-imaging variables most strongly associated are blood pressure, heart rate, and BMI, and the strong delta-expansion indicates that this is a mode strongly reflective of subject variability in the rate of the ongoing aging process. Although these analyses with this data do not allow for strong causal inference, it seems quite plausible that factors such as high blood pressure cause subjects’ brain vasculature and white matter to age in an accelerated, ongoing, way.

### 3.2 Real data - voxel-based deep learning age prediction

We also evaluated delta-expansion from a previous study of brain aging, which had used voxel-level data from just the T1-based structural images, fed into a deep convolutional network (Peng et al., 2021). This approach, which was made openly available, “achieved state-of-the-art performance in UK Biobank data (N= 14,503), with mean absolute error (MAE)= 2.14y in brain age prediction”. This brain-age modelling scenario is therefore very different from the multimodal (structural, functional, connectivity, etc.) IDP-based evaluations described above, although both used UKB data. Even more clearly than with the IDP-based all-in-one results, this brain age modelling showed no evidence of delta-expansion, with *E_f_* = 0.000(*SE* = 0.020) and *B_f_* = 1.000(*SE* = 0.024).

### 3.3 Simulations

From the simulated data, equivalent plots to those shown in Figure 3 were qualitatively identical to the results from real data. For example, the means and ranges of the following measures matched the real data results: the delta-expansion and baseline effects; the correlation *r_xl_* between cross-sectional and longitudinal delta, and the relationship between this and delta-expansion; the correlations between modes and between deltas; the accuracy of age prediction; and the correlations between mode deltas and all-in-one deltas.

Because we know the simulated ground-truth values of some measures (e.g., *B*), we can directly evaluate the accuracy of the estimations of these. We report both correlation and coefficient-of-determination (CoD, 1 − ∑(*B*_est_ − *B*_true_)^2^*/*∑(*B*_true_ − mean(*B*_true_))^2^) between estimated and true values, estimated across the 16 modes within each simulation. CoD is sensitive to errors in scaling and mean value, whereas correlation is not. Table 2 shows the results for several important parameters. We ran the simulation 100 times, and report the mean±std results across these 100 simulations. For these parameters, the inference approach estimates both the measure itself and its expected standard error (uncertainty). Therefore, we report both the expected and actual SE values (averaged over modes); these ideally should match. There is no direct ground-truth definition of these quantities for the all-in-one model, so results are shown for just the modes.

**Table 2:** Accuracy of estimated brain age model parameters compared with ground truth values, for the multiple modes of aging.

| quantity | CoD | r | mean SE (theory) | mean SE (actual) |
| --- | --- | --- | --- | --- |
| $E$ delta-expansion | $0.81 \pm 0.13$ | $0.92 \pm 0.06$ | $0.131 \pm 0.030$ | $0.198 \pm 0.073$ |
| $B$ baseline delta spread | $0.53 \pm 0.24$ | $0.98 \pm 0.01$ | $0.019 \pm 0.001$ | $0.058 \pm 0.005$ |
| $E_f$ fractional expansion | $0.97 \pm 0.02$ | $0.99 \pm 0.01$ | $0.034 \pm 0.002$ | $0.026 \pm 0.005$ |
| $B_f$ fractional baseline | $0.97 \pm 0.03$ | $0.99 \pm 0.01$ | $0.036 \pm 0.002$ | $0.027 \pm 0.005$ |

The theoretically expected (see Appendix A) uncertainties (standard errors) on *E*, *E_f_* and *B_f_* are a reasonable match for the actual SE (root mean square error), whereas the actual error on *B* is larger than predicted, presumably because of the added measurement noise, which inflates apparent baseline.

The correlations of the various models’ deltas with the ground-truth baseline offset and aging rates are shown in Figure 4. For a given simulated mode, each subject has their own baseline offset, and that can be correlated (across subjects) against that mode’s delta values, to indicate how strongly the deltas reflect the baseline effect (recall that the standard deviation of the baseline across subjects corresponds to *B*, which gives the overall strength of the baseline variability for that mode). Similarly, for a given mode, each subject has their own ongoing aging-rate, and that can be correlated against that mode’s delta values, to indicate how strongly the deltas successfully reflect ongoing aging (recall that the standard deviation across subjects of aging rate corresponds to *E*, which gives the overall strength of the delta-expansion for that mode). In most brain aging research, people generally expect delta to reflect rate of ongoing aging, and here the validity (or not) of this assumption is reflected in these delta vs. aging rate correlations.

**Figure 4:**
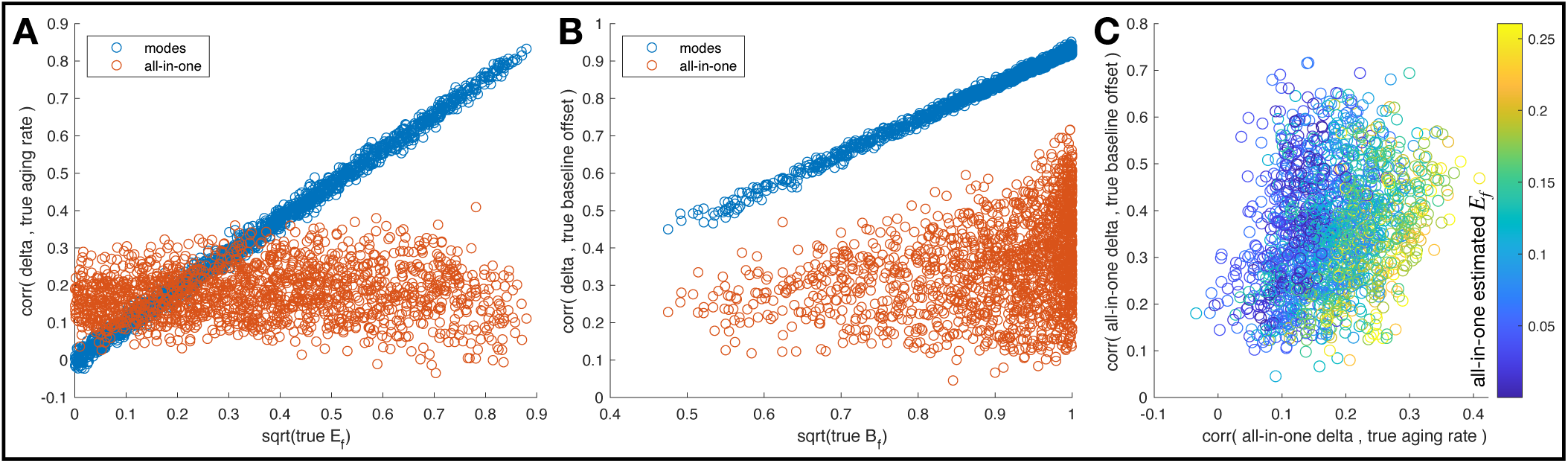
Correlation of different models’ delta estimates with ground-truth subject variations in baseline and aging rate (delta-expansion) effects. The distribution of data points is across 16 modes and 100 simulations.

So, for each mode *j* and subject *i*, the simulation generates a ground-truth baseline *x*_0_*_ij_* and aging rate *β_ij_* which can be compared with the corresponding *δ_xij_* (following Eq. 6). For the all-in-one model, the modes are combined (fitted) to produce just one delta for each subject, meaning that there is no single ground truth baseline or aging rate to compare against. Hence in Figure 4A-B, the all-in-one delta is correlated against the ground truth of each of the 16 modes (resulting in a separate plotted point per mode). Figure 4C focusses on just the all-in-one results, plotting (again by necessity one point per mode) delta-vs-baseline correlation against delta-vs-rate; however, unlike with the *x* axes in 4A-B (which are the true *E_f_* and *B_f_* from the individual modes), here the colour coding reflects the all-in-one estimated delta-expansion. This clearly shows that while the correlation between all-in-one delta and different modes’ aging rates rarely reaches *r* = 0.3 (and is generally much lower), the strength of this correlation is reasonably well predicted by the estimated all-in-one expansion.

For the correlation of delta vs. true aging rate (Figure 4A), the mean (across modes) was 0.16±0.05 for the all-in-one deltas and 0.34±0.06 for the modes. The maximum (across modes) correlation was 0.23±0.05 for the all-in-one deltas and 0.72±0.07 for the modes. The all-in-one deltas cannot of course be expected to correlate with the subject aging rates of individual imaging features (or modes) unless there is reasonable correlation between the modes; because we have simulated correlation between modes here in a way that matches the real data, these correlations for the all-in-one deltas (0.16) are poor but not zero. The maximum delta vs. aging-rate correlation for the individual modes is for those modes that have the greatest delta-expansion (and hence valid ongoing aging effects); as a result this is considerably higher than the maximum correlation seen with the all-in-one model. For the modes, it is clear that the correlations of deltas with the true aging rate is proportional to (indeed virtually equal to) the square root of the true fractional variance explained by the delta-expansion.

We now return to the potential modelling problem of non-zero correlations between true baseline and delta-expansion, which might occur if delta-expansion had been happening before the time period of the study. The correlation between cross-sectional and longitudinal delta estimates might be used to identify the presence of correlations between baseline and expansion effects, but the presence of measurement noise reduces the former to the extent that it is not a good predictor of the latter. Therefore we reran the simulations, this time applying the “worst-case scenario” of perfect correlation between true baseline and expansion. We then estimated the error in estimation of *E_f_* and *B_f_* . The results are shown in Figure 5.

**Figure 5:**
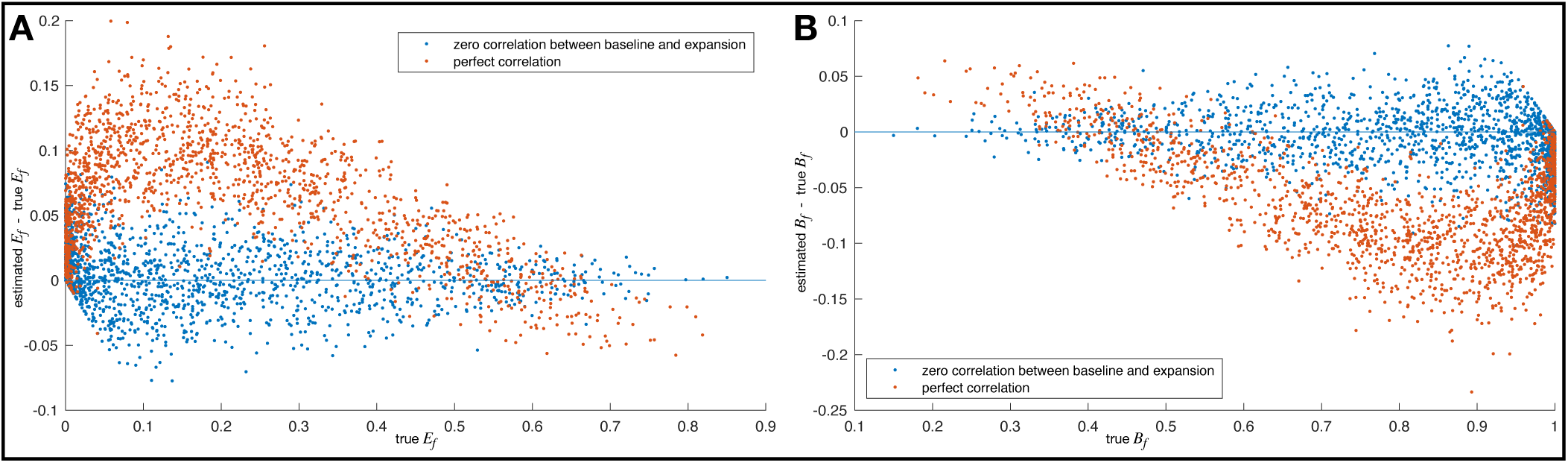
Error in estimation of *E_f_* and *B_f_* when true baseline and expansion are correlated. The distribution of data points is across 16 modes and 100 simulations.

The blue data points show the estimation accuracy when true baseline and expansion are uncorrelated. There is no obvious bias, and the spread of errors is consistent with the standard errors reported in Table 2 (i.e., approximately 0.03). The orange data points show the estimation accuracy when true baseline and expansion are perfectly correlated - an unlikely worst-case scenario. However, even in this scenario, the biases on *E_f_* and *B_f_* are quite small, rarely exceeding 0.15; furthermore, for true *E_f_ >* 0.4, the bias almost never rises above 0.05. Overall, therefore, it does not appear that such correlations would present a significant problem for the estimation and interpretation of these model parameters.

## 4 Discussion

“All-in-one” models of aging in general do not strongly inform us about between-subject variations in the rate of ongoing aging, and do not do a good job of reflecting the multiple biological processes underlying aging. Part of the underlying problem is illustrated in Figure 6, which shows a simplified causal graph for aging processes in the brain. Some aging pathways involve interactions between aging factors, for example, in aging mode 9, it is likely that blood pressure is influencing the rate of aging, as seen in the increases in white matter lesion volumes, whereas in mode 1 our results suggest that there is no factor modulating the effect of aging. There is nothing in this generative model of brain imaging features that “points towards” using an all-in-one model to understand brain aging.

**Figure 6:**
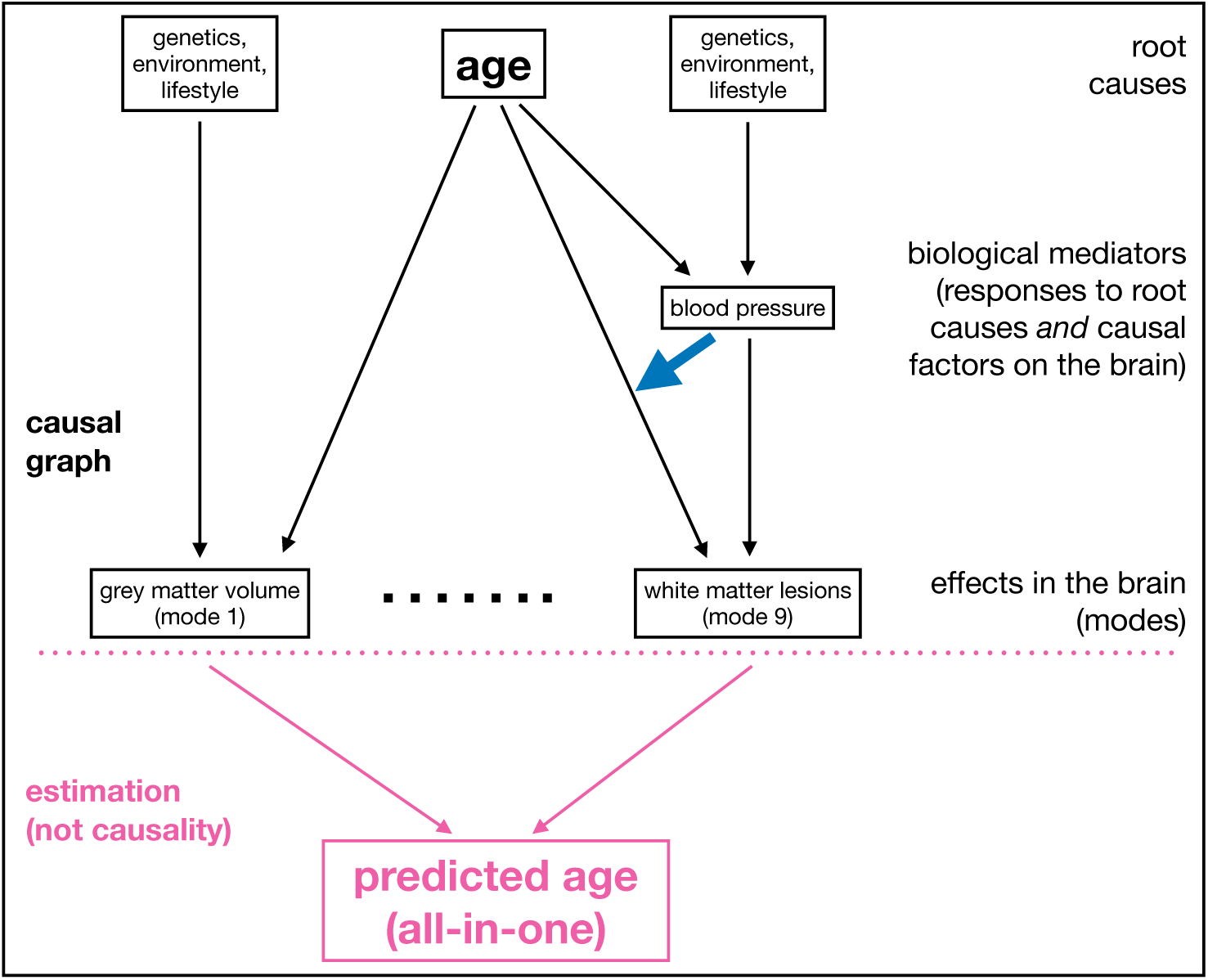
Simplified causal graph for aging processes in the brain.

Related to this conceptual distinction between all-in-one modelling vs. generative models for distinct modes of aging, we can revisit the all-in-one model using a simplified re-formulation, in order to add further qualitative interpretation of the model’s fitting process. The generative models for modes *x*_1_*, x*_2_*, …* are:

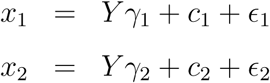

where *Y* is age and *γ_j_* is the population-average aging-rate for mode *j*. *c_j_* is a mode-specific combination of other causes on a brain mode (for example, smoking or blood pressure), which is typically present but not known a priori; *c_j_* will be independent of age, as any age-related component will get wrapped into the first term, altering *γ*. Finally there is a random error term *ɛ_j_* that is uncorrelated between the different modes, e.g., reflecting imaging measurement noise.

The all-in-one model estimates the linear combination of image features *x_j_* that optimally predicts age:

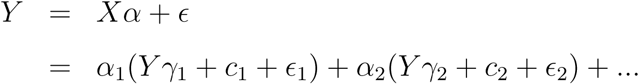

Hence the all-in-one model will be seeking to optimise age prediction by averaging across the multiple noise sources *ɛ_j_* while finding the optimal set of weights to combine the modes together (those weights being partially driven by *γ_j_* and varying *ɛ_j_* noise levels). However, crucially, the all-in-one-model must also optimise regression coefficients such that the effect of other causal factors *c_j_* is minimally detrimental to the age prediction. This may involve adjusting regression coefficients such that (unknown but present) nuisance causal factors *c_j_* cancel each other out, where they correlate across modes.

Hence, again, this view on these models undermines the biological interpretability of the all-in-one model. The extent to which different modes contribute to the all-in-one model varies highly (including regression coefficients going negative), and is not driven simply by individual modes’ age correlation (as described above, and empirically, see the difference between the first and third plots in Figure 3F), but by the correlation structure between the modes. This is consistent with the problem that the natural generative model has age causing each distinct mode of brain aging (in that direction, and in different ways for each biological mode), and not the other way round. Combining across multiple modes may achieve the best-possible all-in-one age prediction, but does not provide for biological interpretability.

It is encouraging that we did find positive correlation *r_xl_* between cross-sectional and longitudinal deltas; this was the case even for the all-in-one model, which was small (*r_xl_* = 0.05) but statistically significant. We did however find that several IDP-preprocessing steps were important in order for the all-in-one *r_xl_* to be estimated as positive, and it is worth noting these here. Originally, we trained the all-in-one brain age regression model using all subjects (tens of thousands), and not just those with two timepoints; when applied to those subjects with two timepoints, we found a negative *r_xl_*, and realised that there was a small healthy bias in those subjects who have returned for a second scan, in the sense that their average deltas were negative (whereas by definition in any given training dataset the average delta is zero). Retraining age prediction in a simpler, unbiased way, using just the two-timepoint subjects, eliminated this bias and the resulting negative *r_xl_*. Secondly, we found that it is important to include the imaging confounds when adjusting for brain age bias (S. Smith et al., 2019), and not just in IDP pre-processing, to avoid age-correlated confounds from implicitly re-appearing, and affecting measures like *r_xl_*. Similarly, we found that in some cases, IDP quantile normalisation (often applied by default in IDP-preprocessing), which reduces extreme IDP values, was skewing some longitudinal delta estimation. Finally, it is important to correlate longitudinal delta with a cross-sectional estimate formed from the average of the two timepoints, because using just the first or second timepoint on its own brings a strong noise-correlation bias.

One subtlety of brain age modelling that may be under-appreciated is the importance of the population-average fit to the interpretation of subject-level deltas. For example, in scenarios where age prediction is poor, the “delta” values are virtually identical to the imaging features fed in, because there is little age dependence to subtract off. Where this is the case, it seems that it makes little sense to interpret delta as relating to aging at all. However, with the expansion model, even in the absence of accurate age prediction (i.e., even when the population average age curve does not have strong age dependence), one could still be confident that large estimated delta-expansion is indicative of meaningful brain age modelling. Conversely, if age prediction accuracy is very high, but there is no evidence of significant delta-expansion, it is unlikely to be reasonable to interpret brain age delta values as indicating accelerated or slowed ongoing brain aging.

One might ask “surely when brain age prediction is highly accurate, that must by definition mean that we have found a meaningful signature of aging in the imaging data - in which case, the deviation of that brain age from actual age must also mean something?”. This perspective is intuitive, but conflates the effects of what one might term as “healthy aging”, where individuals age at the same rate, and “unhealthy aging”, where some individuals age faster due to pathological processes. For example, the strongest mode of brain aging primarily reflects total grey matter volume, which indeed is known to decrease strongly in aging; however, this simple brain measure also of course varies a lot across the population, varying with birth weight, sex, overall body size, and many other factors occurring during the first years of life (Vidal-Pineiro et al., 2021). It is not incompatible therefore to see a brain measure showing major baseline variability as well as strong cross-sectional (and even longitudinal) variation with aging, and yet not necessarily predominantly reflecting subject-specific aging rates in older life.

How useful are these models in the presence of pathology? Some pathologies (e.g., some dementias) may look spatially similar to normal aging, but with even greater acceleration, and the models, even when fully pre-trained on a population of healthy aging, may be immediately useful for helping quantitate the state (and/or rate of progression) of the disease. Other pathologies (e.g., stroke) may show gross structural effects that are not well captured by the patterns seen in normal aging, even when characterised as multiple aging modes, and it will be important to apply clinically-informed interpretation in such cases. Similarly, how relevant are these models to other life stages, for example during fetal and infant periods? We would expect the models and interpretations proposed here to be relevant to datasets covering other age ranges (and organs, and species), but would not expect specific model training (the learning of the population average age curves) to translate well between highly different datasets - that would need to be relearned in general.

In conclusion, we believe that reasonable interpretation of brain age delta requires an understanding of the extent to which delta is reflecting between-subject baseline variations (meaning that all subjects are actually aging at the same rate), vs. between-subject variations in the rate of ongoing aging. In addition, it seems likely that most “all-in-one” delta estimates largely reflect baseline effects, whereas the estimation of multiple biologically distinct modes of aging allows to find a variety of baseline-vs-expansion scenarios, including some that do show strong between-subject variability in ongoing aging rates.

## Data and Code Availability

Real data (e.g., IDPs and confounds) is available following an application process to UKB (this work used UKB data accessed via data access application 8107). All new matlab code for processing real data and generating simulations is openly available from https://tinyurl.com/baneraging (this will be updated to being a Zenodo/etc download prior to finalising publication).

## Author Contributions

All authors contributed to study design, data analysis, and writing.

## Acknowledgements

We are grateful to UK Biobank and all UK Biobank participants for their time and effort in creating this rich imaging resource. We are also grateful to Han Peng for providing the deep-learning-based age predictions.

## A Estimation of delta baseline and expansion

Consider brain age delta for subject *i* and feature *j* (*δ_X,ij_*) – in this appendix we denote this *δ_X,i_*, suppressing feature index as all modelling here will be applied to each feature separately. We assume that *δ_X,i_* is mean zero but is the sum of two random components

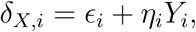

where *ɛ_i_* and *η_i_* are independent random variables with Var(*ɛ_i_*) = *σ*^2^ and Var(*η_i_*) = *τ* ^2^; the first component is age-invariant, and the second expresses a monotypic dependence on age (as above, we assume ages have been rescaled to [0, 1]).

The variance of brain age delta is then

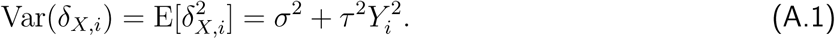

We note that this model makes a very specific assumption about the shape of the heterogeneity of variance, with variance being minimal at the lowest age and increasing as a multiple of *Y_i_*^2^.

Estimation of *σ*^2^ and *τ* ^2^ from *δ_X,i_* proceeds as follows. Assuming a Gaussian distribution, the log likelihood of the *N* observations is

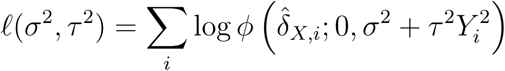

where *ϕ*(*t*; ·, ·) is the PDF of a Normal distribution evaluated at *t* with a given mean and variance. Any optimizer can be used to maximise *ℓ*(*σ*^2^*, τ* ^2^), but to obtain the most robust results we use a succession of methods:

1. A methods of moments estimate is first found, obtained by regressing *δ_X,i_*^2^ on *Y_i_*^2^, such that the intercept estimates *σ*^2^ and the slope *τ* ^2^ (see Eqn. (A.1)); any negative estimates are set to zero.
2. A robust simplex method (Matlab’s ‘fminsearch‘) is used to get a better estimate, using the previous estimate as a starting value.
3. A line search method (Matlab’s ‘fminunc‘) is used, using the previous estimate as starting value, to get a more precise estimate and the Hessian at the optimised value.

To simplify the implementation, an unconstrained optimization is used with log likelihood parameterized as *ℓ*(*σ, τ*), and absolute values are taken after optimization.

The Hessian is the observed Fisher information, *I*(*σ, τ*), and its inverse gives asymptotic variance-covariance variance of [*σ*^; *τ*^]; denote the square-root diagonals of *I*^−1^(*σ*^*, τ*^) as SE(*σ*^) and SE(*τ*^), respectively. The total variance of *δ_Xi_* is a sum of two components

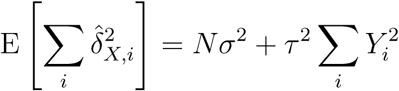

and thus the proportion of variance attributable to ‘growing’ variance is

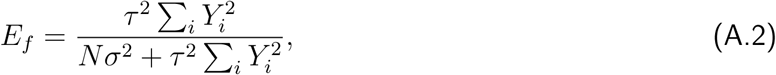

though in practice we estimate the denominator with the total data sums of squares, ∑*_i_δ_X,i_*^2^. We found the delta-method variance, Var(*τ*^^2^) ≈ (2*τ*^)^2^(SE(*τ*^))^2^, to be quite inaccurate, and so instead compute SE(*τ*^^2^) from *I*(*σ*^2^*, τ* ^2^) using *ℓ*(*σ*^2^*, τ* ^2^) evaluated at the optimised (squared) estimates.

To consider the possibility of “contracting” delta, we also repeat the above modelling with *Y* replaced by 1 − *Y*, and use the version with the larger optimised log likelihood value.

## B Relationship between different models’ age predictions and deltas

The relationship between the all-in-one age prediction and the individual modes of aging *X_j_* is straight-forward. From the all-in-one model, *Y*_brain_ = *Xα* = ∑*_j_α_j_X_j_*, and hence the all-in-one regression parameters *α* fully define the relationship.

The relationship between all-in-one deltas and individual modes’ deltas is found:

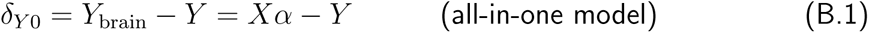

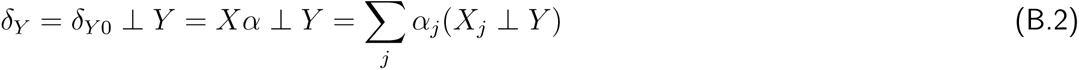

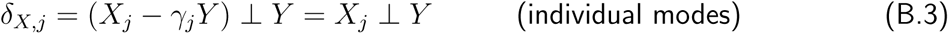

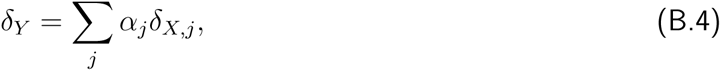

hence giving a simple relationship between deltas. The correlation between deltas is therefore given by:

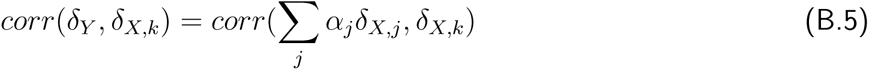

Therefore, because in practice we find that most modes’ deltas correlate positively with each other, and because most of the *α_j_* values are positive in practice, we would expect that all modes’ deltas to correlate positively with the all-in-one deltas.

## References

1. Alfaro-Almagro, F., Jenkinson, M., Bangerter, N., Andersson, J., Griffanti, L., Douaud, G., Sotiropoulos, S., Jbabdi, S., Hernandez-Fernandez, M., Valee, E., Vidaurre, D., Webster, M., McCarthy, P., Rorden, C., Daducci, A., Alexander, D., Zhang, H., Dragonu, I., Matthews, P., … Smith, S. (2018). Image processing and quality control for the first 10,000 brain imaging datasets from UK Biobank. NeuroImage, 166, 400–424.

2. Cole, J. H., & Franke, K. (2017). Predicting age using neuroimaging: Innovative brain ageing biomarkers. Trends in Neurosciences, 40, 681–690.

3. Cole, J. H., Poudel, R. P., Tsagkrasoulis, D., Caan, M. W., Steves, C., Spector, T. D., & Montana, G. (2017). Predicting brain age with deep learning from raw imaging data results in a reliable and heritable biomarker. NeuroImage, 163, 115–124.

4. Cordes, C., Baum, T., Dieckmeyer, M., Ruschke, S., Diefenbach, M. N., Hauner, H., Kirschke, J. S., & Karampinos, D. C. (2016). MR-Based Assessment of Bone Marrow Fat in Osteoporosis, Diabetes, and Obesity. Front Endocrinol (Lausanne), 7, 74.

5. Franke, K., Ziegler, G., Klöppel, S., Gaser, C., & ADNI. (2010). Estimating the age of healthy subjects from T1-weighted MRI scans using kernel methods: Exploring the influence of various parameters. NeuroImage, 50, 883–892.

6. Hastie, T., & Tibshirani, R. (2017). *Generalized additive models*. Routledge, Taylor & Francis. 10.1201/9780203753781

7. Hyvärinen, A. (1999). Fast and robust fixed-point algorithms for independent component analysis. IEEE Transactions on Neural Networks, 10 (3), 626–634.

8. Kessler, D., Angstadt, M., & Sripada, C. (2016). Growth Charting of Brain Connectivity Networks and the Identification of Attention Impairment in Youth. JAMA Psychiatry, 73 (5), 481–489.

9. Loskutova, N., Honea, R. A., Vidoni, E. D., Brooks, W. M., & Burns, J. M. (2009). Bone density and brain atrophy in early Alzheimer’s disease. J. Alzheimers Dis., 18 (4), 777–785.

10. Miller, K., Alfaro-Almagro, F., Bangerter, N., Thomas, D., Yacoub, E., Xu, J., Bartsch, A., Jbabdi, S., Sotiropoulos, S., Andersson, J., Griffanti, L., Douaud, G., Okell, T., Weale, P., Dragonu, I., Garratt, S., Hudson, S., Collins, R., Jenkinson, M., … Smith, S. (2016). Multimodal population brain imaging in the UK Biobank prospective epidemiological study. Nature Neuroscience, 19, 1523–1536.

11. Moqri, M., Herzog, C., Poganik, J., Justice, J., et al. (2023). Biomarkers of aging for the identification and evaluation of longevity interventions. Cell, 186 (18), 3758–3775.

12. Peng, H., Gong, W., Beckmann, C. F., Vedaldi, A., & Smith, S. M. (2021). Accurate brain age prediction with lightweight deep neural networks. Medical Image Analysis, 68. 10.1016/j.media.2020.101871

13. Smith, S., Vidaurre, D., Alfaro-Almagro, F., Nichols, T., & Miller, K. (2019). Estimation of brain age delta from brain imaging. NeuroImage, 200, 528–539.

14. Smith, S. M., Elliott, L. T., Alfaro-Almagro, F., McCarthy, P., Nichols, T. E., Douaud, G., & Miller, K. L. (2020). Brain aging comprises many modes of structural and functional change with distinct genetic and biophysical associations. eLife, 9, e52677. 10.7554/eLife.52677

15. Vidal-Pineiro, D., Wang, Y., Krogsrud, S. K., Amlien, I. K., Baaré, W. F., Bartres-Faz, D., Bertram, L., Brandmaier, A. M., Drevon, C. A., Düzel, S., Ebmeier, K., Henson, R. N., Junqúe, C., Kievit, R. A., Kühn, S., Leonardsen, E., Lindenberger, U., Madsen, K. S., Magnussen, F., … Fjell, A. (2021). Individual variations in ‘brain age’ relate to early-life factors more than to longitudinal brain change. eLife, 10, e69995. 10.7554/eLife.69995

16. Vinke, E. J., de Groot, M., Venkatraghavan, V., Klein, S., Niessen, W. J., Ikram, M. A., & Vernooij, M. W. (2018). Trajectories of imaging markers in brain aging: the Rotterdam Study. Neurobiol. Aging, 71, 32–40.

